# A simple model for the evolution of temperature-dependent sex determination explains the temperature sensitivity of embryonic mortality in imperiled reptiles

**DOI:** 10.1101/791202

**Authors:** Lauren Lawson, Njal Rollinson

## Abstract

A common reptile conservation strategy involves artificial incubation of embryos and release of hatchlings or juveniles into wild populations. Temperature-dependent sex determination (TSD) occurs in most chelonians, permitting conservation managers to bias sex ratios towards females by incubating embryos at high temperatures, ultimately allowing the introduction of more egg-bearing individuals into populations. Here, we revisit classic sex allocation theory and hypothesize that TSD evolved in some reptile groups (specifically, chelonians and crocodilians) because male fitness is more sensitive to condition (general health, vigor) than female fitness. It follows that males benefit more than females from incubation environments that confer high-quality phenotypes, and hence high-condition individuals. We predict that female-producing temperatures, which comprise relatively high incubation temperatures in chelonians and crocodilians, are relatively stressful for embryos and subsequent life stages. We synthesize data from 28 studies to investigate how constant temperature incubation affects embryonic mortality in chelonians with TSD. We find several lines of evidence suggesting that female-producing temperatures, especially relatively warm temperatures, are more stressful than male-producing temperatures, and we find some evidence that pivotal temperatures (TPiv, the temperature that produces a 1:1 sex ratio) exhibit a correlated evolution with embryonic thermal tolerance. If patterns of temperature-sensitive embryonic mortality are also indicative of chronic thermal stress that occurs post hatching, then conservation programs may benefit from incubating eggs close to species-specific TPivs, thus avoiding high-temperature incubation. Indeed, our models predict that, on average, a sex ratio of more than 75% females can generally be achieved by incubating eggs only 1°C above TPiv. Of equal importance, we provide insight into the enigmatic evolution of TSD in chelonians, by providing support to the hypothesis that TSD evolution is related to the quality of the phenotype conferred by incubation temperature, with males produced in high-quality incubation environments.

**Lay summary:** We analyze data on embryonic mortality under constant-temperature incubation for 15 species of chelonians with temperature-dependent sex determination (TSD). Mortality is lowest near species-specific pivotal temperatures (Tpiv) but increases rapidly above TPiv, consistent with a theory that explains the adaptive significance of TSD. Conservation managers should incubate embryos near TPiv.

## Introduction

Despite recent and widespread interest in reptile conservation (Roll *et al.*, 2017), reptile populations are declining globally (Todd *et al.*, 2010). Turtles, for example, are among the most imperiled group of vertebrates in the world (Rhodin *et al.*, 2018; Gibbons and Lovich, 2019; Stanford *et al.*, 2020). Natural rates of replacement and population growth are low in most turtle species, and because of high juvenile mortality, slow life histories (e.g., Kondo, Morimoto, Sato, & Suganuma, 2017), and low genetic diversity (Romiguier *et al.*, 2014), populations are unable to adapt to environmental change on the time scale of anthropogenic impacts (Hawkes *et al.*, 2009). Elevated adult mortality of turtles arising from road collisions (Steen and Gibbs, 2004), fishing gear entrapment (Lewison *et al.*, 2004; Bolten *et al.*, 2011), predation (Bolten *et al.*, 2011), and direct consumption as a food source (Conway-Gómez, 2007; Hancock *et al.*, 2017), has therefore resulted in dramatic population declines. At present, 56.3% of data sufficient species (51.9% of all recognized species) are considered critically endangered, endangered, or vulnerable by the International Union for Conservation of Nature (Rhodin *et al.*, 2018).

Common turtle conservation initiatives include protecting nesting turtles from natural predation and poachers (Eckert *et al.*, 1999), rehabilitation programs at trauma centers (Feck and Hamann, 2013), education initiatives (Hassan *et al.*, 2017), and captive breeding (Bowkett, 2009). Another common initiative undertaken by zoos, parks, conservation authorities, and even private individuals, involves incubation of reptile eggs in an ex-situ setting (Eckert *et al.*, 1999), followed by the release of hatchlings or juveniles. Growth of wild populations may then be enhanced by protecting eggs and embryos, which reduces embryonic and hatchling mortality, and/or rearing individuals to larger body sizes before release (Carstairs *et al.*, 2019; Tetzlaff *et al.*, 2019), which may increase survival rates in the wild (Rollinson and Rowe, 2015).

Artificial incubation is an integral part of many conservation initiatives for reptiles. Artificial incubation typically follows the removal of eggs from the wild (e.g., natural nests, or gravid females that died on a road), or egg production after captive breeding. Embryos are placed in incubators, hatched under a specified incubation regime, then released into the wild (Páez *et al.*, 2015). Although the physical incubation of reptilian embryos may seem to represent a small or insignificant portion of conservation programs, it is widely recognized incubation temperature has a profound influence on morphology, performance, and fitness components (Bobyn and Brooks, 1994a, 1994b; Booth, 2006; Noble *et al.*, 2018b). Further, some lizards, all crocodilians, the tuatara, and most turtles exhibit temperature-dependent sex determination (TSD), where sex is permanently affected by incubation temperature (Valenzuela and Lance, 2004). The evolution and maintenance of TSD may in fact be related to the effect of incubation temperature on fitness components (West, 2009), and as such, exploring artificial incubation regimes for reptiles through the lens of TSD evolution may provide insight into best practices for conservation programs.

In reptiles with TSD, the incubation temperatures experienced by an embryo during the thermosensitive period influences sex, where the thermosensitive period comprises specific anatomical stages that occur roughly during the middle third of embryonic development (Yntema, 1968, 1979; Girondot *et al.*, 2018). Under constant temperature, the temperature-sex reaction norm is known to take three forms (reviewed in Valenzuela and Lance, 2004). The FMF pattern occurs when males are produced at intermediate temperatures and females are produced at extreme temperatures. FMF is hypothesized to be ancestral, and it is found in some turtles, lizards, and all crocodilians. The MF pattern occurs when males are produced at cool temperatures and females at hot temperatures. Although turtles are the only taxon to exhibit MF, turtles happen to comprise the majority of extant TSD species, and therefore MF is the most common pattern of TSD. Finally, the FM pattern is when females are produced at cool temperatures and males at hot temperatures; but this pattern is rare and found only in a few lizards and the tuatara. Evolutionary explanations for TSD in lizards and the tuatara (FM, and FMF) may be different than evolutionary explanations for TSD in turtles and crocodilians (MF, and FMF) (e.g., Janzen and Phillips 2006). For instance, squamates are a sister group to the tuatara (Rest *et al.*, 2003), whereas turtles are sister to the Archosaurs, which includes crocodilians (Crawford *et al.*, 2012), underlining that fundamentally different patterns of TSD are found in divergent evolutionary lineages.

In general, the evolution of TSD is hypothesized to arise by virtue of a sex-by-temperature interaction for fitness (Charnov and Bull 1977). In short-lived squamates with rapid maturation, the FM pattern of TSD may represent an adaptation to ensure females hatch under the cool conditions that occur relatively early in the season, as to maximize time for growth and achieve high fecundity during their short lives (Pen *et al.*, 2010, see also Conover, 1984). In contrast, turtles and crocodilians mature late and are very long lived, so TSD may be an adaptation to an effect of incubation temperature on adult fitness that occurs over their relatively longer lives (Figure 1). Specifically, males generally experience competition for mates, and a small proportion of males often capture a large portion of mating opportunities (Bateman 1948), with males in high “condition” (i.e., vigorous males in good health) securing most mates (Rowe and Houle, 1996). Male fitness is therefore more sensitive to condition than female fitness (Trivers and Willard, 1983), such that TSD evolves in age- and size-structured populations to ensure that males are produced under temperatures that represent a high-quality incubation environment, conferring relatively high post-hatching condition, ultimately enhancing male fitness under strong sexual selection (Trivers and Willard, 1973; Deeming and Ferguson, 1989; West, 2009). This hypothesis is an extension of Trivers-Willard (1973).

**Figure 1:**
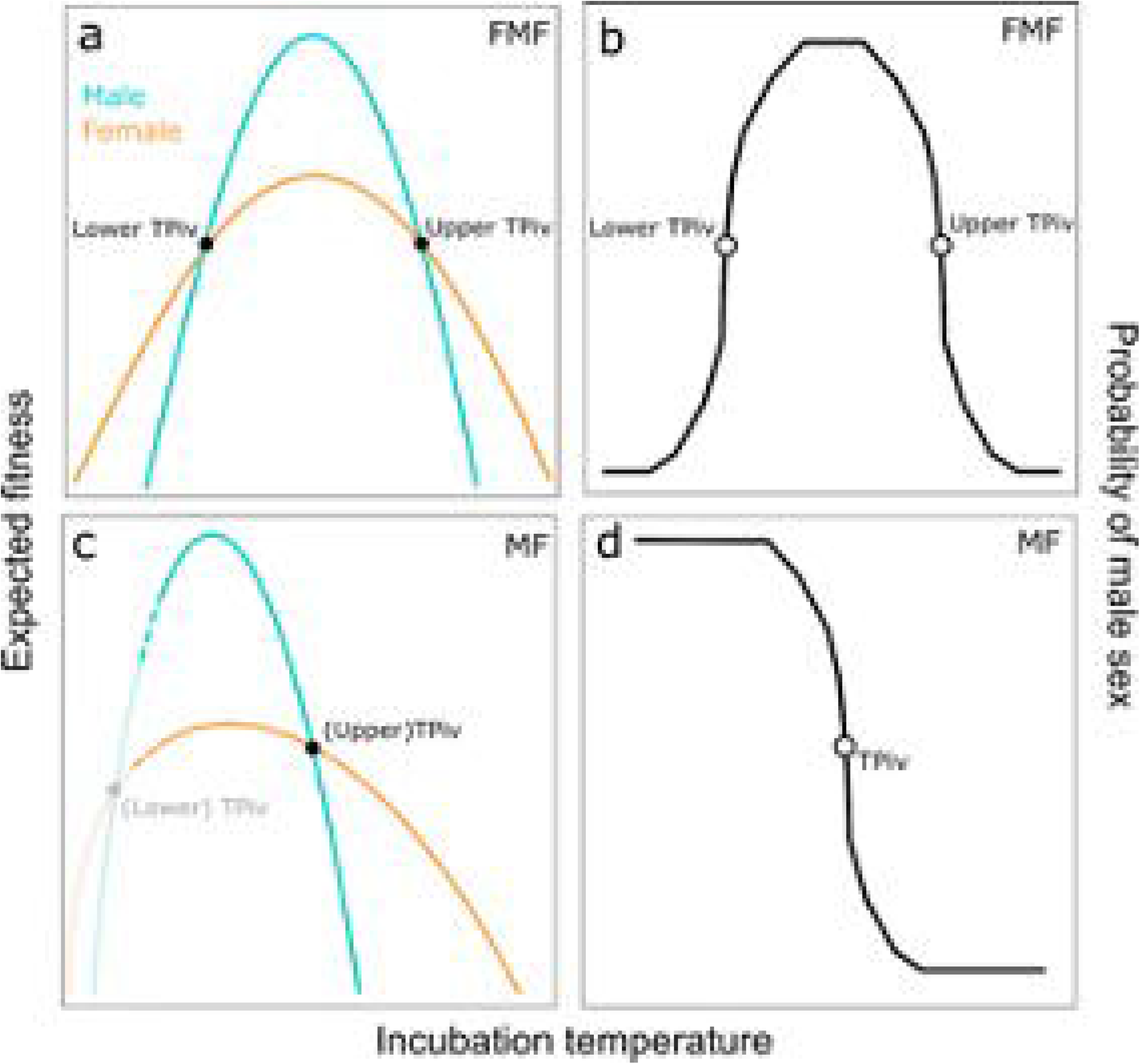
Conceptual model for the evolution of TSD in turtles (and crocodilians), with explicit links between pivotal temperatures (TPiv), expected fitness, and reaction norms for sex. Types of TSD are (a) an “FMF” pattern (or Type II TSD) or (c) an “MF” pattern (or Type Ia TSD). An optimal incubation temperature produces high-quality phenotypes as in FMF species (panel a), or lower temperatures have low developmental leverage and have little influence on sex as in MF species (panel c). In either case, males should be produced under the conditions that maximize adult fitness for both sexes, as males stand to gain more fitness from good incubation environments, and to lose more fitness from bad incubation environments. This is equivalent to a sex-by-environment interaction for fitness, which gives rise to (b) the FMF reaction norm, or a special case of the FMF reaction norm, which is (d) the MF reaction norm. In panel (c) the lower TPiv (light grey) indicates females can be produced under extremely low temperatures in MF species.

The FMF pattern can easily be reconciled with our hypothesis, where females are produced in extreme environments (i.e., hot and cold incubation environments), and males at intermediate, or “optimal”, temperatures (Fig 1a). The MF pattern can, however, also be reconciled with our hypothesis: because sex is determined by the amount of development that occurs above vs below TPiv (Georges *et al.*, 1994), lower temperatures must have relatively little effect on sexual differentiation as they have little developmental leverage (Rollinson *et al.*, 2018; Massey *et al.*, 2019), and therefore natural selection to maintain the lower TPiv is weak in FMF species; this ultimately gives rise to MF species (Fig 1). Thus, a critical prediction of our hypothesis is that fitness should decline with temperature at a relatively faster rate above TPiv vs below the TPiv for MF species, and above vs below the upper TPiv for FMF species. The reasoning is that male-producing temperatures should reflect the peak of the performance curve, where a unit change in temperature has little influence on fitness, whereas female-producing temperatures should represent the shoulder of the curve, where fitness changes rapidly with temperature (Fig. 1). Therefore, performance metrics should be relatively invariant with respect to temperature in the range of male-producing temperatures (i.e, slightly cooler than TPiv), although male-producing temperatures that are far cooler than TPiv may result in a decrease in performance.

We have outlined a theoretical ground to suggest that incubation temperature has predictable and long-term effects on phenotypes and fitness components, especially in turtles and crocodilians. Yet, incubation regimes in conservation programs for these taxa are broadly unstandardized. For instance, some head-starting programs allow natural egg incubation (Bona *et al.*, 2012), some allow temperature to fluctuate with above-ground ambient temperature (Tetzlaff *et al.*, 2019), and others incubate different sets of eggs at different constant temperatures (Carstairs *et al.*, 2019). A broad and theoretically informed exploration of how incubation temperature affects fitness in embryos and hatchlings may therefore be useful in developing best practices for artificial incubation in conservation programs.

Our goal is to better understand the evolution of TSD, and to inform best practices for artificial incubation programs geared towards the conservation of reptiles, particularly for chelonians. In the present study, we synthesize existing data to explore the fitness consequences of artificial incubation regimes through the lens of TSD theory. We focus on patterns of temperature-sensitive embryonic mortality in chelonians, as chelonians are relatively data rich, and mortality is both a fitness component and is commonly measured. We generalize our mortality findings by centering study-specific incubation temperatures with respect to species-specific pivotal temperatures (TPiv), which is the constant temperature at which the sex ratio is 1:1 (Bull, 1980; Ewert *et al.*, 1994). We focus on a simple explanation for TSD that is specific to chelonians and crocodilians (Figure 1), which is that males are produced in incubation environments that confer high condition (Charnov and Bull, 1977; Deeming and Ferguson, 1989; West, 2009). Several predictions arise from this hypothesis. The first is that embryonic thermal tolerance exhibits a positive correlated evolution with TPiv, where thermal tolerance and TPiv values evolve together. This pattern occurs because embryonic physiology is locally adapted to incubation regimes (Ewert 1985), and TPiv evolution ensures that extreme temperatures within local regimes produce female hatchlings, consistent with the adaptive explanation for TSD. Therefore, we predict that (1) embryonic mortality will be best explained by the difference between observed incubation temperature and population-specific TPivs, compared to the difference between observed incubation temperature and mean incubation temperature, or simply the observation incubation temperature. We also predict that (2) mortality will be positively associated with the difference between incubation temperature and TPiv. Next, under the adaptive explanation for TSD explored herein, temperature affects individual quality and males are produced in high-quality thermal environments, but the association between sex and individual quality arises through temperature. We therefore predict that (3) sex ratio (percent male) will be negatively associated with embryonic mortality, but after accounting for temperature, mortality will not be associated with sex. Finally, if TPiv evolves to delineate high-quality thermal environments (for male production), from low quality thermal environments (for female production), then (4) embryonic mortality in chelonians should increase with temperature more rapidly above the TPiv, compared to below the TPiv. Mechanistically, fitness should depreciate rapidly above the TPiv because TPiv evolves to be closer to the upper end of thermal tolerance, rather than the lower end, and reaction rates for most or all processes changes more rapidly near the upper tolerance limit (Sharpe and DeMichele, 1977; Kingsolver, 2009).

## Methods

### Data collection

Data was collected from previously published studies collated in the Reptile Development Database (Noble *et al.*, 2018a). Only chelonians were considered in this study, as they are of extreme conservation concern (Rhodin *et al.*, 2018; Gibbons and Lovich, 2019; Stanford *et al.*, 2020), they represent the overwhelming majority of species in the world that exhibit TSD, they feature a pattern of TSD different from squamates and the tuatara (see below), and very limited data was available on crocodilians, especially for TPivs. Thus, studies from the Reptile Development Database were included in the present study if they featured (1) a turtle species with TSD (Sabath *et al.*, 2016), (2) artificial incubation under constant temperature, (3) embryonic mortality (or survival) specific to a temperature treatment, and (4) the number of embryos comprising the sample. Data collected from studies fitting inclusion criteria included species, incubation temperature, TPiv, percent mortality per incubation regime, number of eggs incubated, and geographic location of egg collection. Exclusion factors included genotypic sex determined species (GSD), in situ incubation, or fluctuating temperature trials. In studies including both TSD and GSD species, and studies including both constant and fluctuating incubation trials, only TSD and constant trial results were extracted. Trials involving temperature shifts were also excluded. Several studies report mortality results in figures and do not specify results textually; in these cases we extracted data from figures using WebPlotDigitzer (Rohatgi, 2019). We also extracted all available information on substrate moisture; however, moisture was only measured directly in ten studies. Notably, all studies either reported moisture values or maintained moisture at approximately constant levels across the incubation regime; further, the few studies that manipulated moisture and temperature did so in a factorial manner. It is therefore unlikely that moisture was confounded with temperature in the present study.

Ultimately, 28 studies in the Reptile Development Database fit inclusion criteria for the current study (Supplementary Table 1). In total, 199 data points were collected on a total of 15 different species in 6 different families of turtle, representing 43% of known taxonomic families.

### Pivotal temperatures

Some papers that were used in the present study did not explicitly state the population-specific TPiv(s). These values were supplemented with TPivs from other papers studying the exact same population, or the most proximal population of the same species with a TPiv estimate, as TPiv can vary geographically and among species (Bull, 1980; Ewert *et al.*, 1994; Carter *et al.*, 2019). To accomplish this, we performed literature searches to find TPivs of species using the species name, ‘pivotal temperature’, and ‘sex ratios’ as search terms. Species-specific TPivs were found, and the closest geographically available population TPiv found was used as the TPiv. In total, thirteen studies required supplementation of TPiv from previously published literature (Supplementary Table 1).

There was only one species with FMF TSD (Pattern II), the snapping turtle (*Chelydra serpentina*) in which both cool and hot temperatures produce females (Figure 1). For this species, we use the upper TPiv in our analyses, since high temperature incubation is the focus of this study; however, we perform all analyses with and without this species to ensure patterns are robust regardless of the type of TSD.

### Data Analysis

We reasoned that the relationship between temperature and embryonic mortality is likely non-linear (e.g., Amarasekare and Savage, 2012). We therefore allowed flexibility in fixed effects in all models (described below) by using basis splines within the package *splines* (R Core Team, 2020). We chose basis splines over natural splines as basis splines are not constrained to linearity at the tails, which is important as the tails of our functions are associated with few data points, and basis splines therefore emphasize the uncertainty, rather than minimize it. Except where noted, embryonic mortality was always the dependent variable, and it was weighted by sample size of embryos in each temperature treatment. Given that many species were represented by only one study, we chose to use only StudyID as a random intercept. We explored the possibility of including basis splines as random slopes within StudyID, but these models generally exhibited signatures of overfitting, and results were broadly robust whether or not random slopes were included. Having selected our lone random effect to be applied in all models, our models were subsequently fit using maximum likelihood. Model comparisons outlined below were performed using second order Akaike Information Criterion (AIC_c_) and Akaike weights (*w*_i_) within the package ‘AICmodavg’ in R (Fabozzi *et al.*, 2014; Mazzerole, 2019); basis spline regression was applied to fixed effects using the *bs* function in *splines* and *lmer* in lme4 (Bates *et al.*, 2015).

We expect that TPiv exhibits a correlated evolution with embryonic thermal tolerance. To address this expectation, we generated three plausible hypotheses for how incubation temperature affects mortality, and we expressed these hypotheses as statistical models (Burnham and Anderson 2002). The first and simplest hypothesis is that mortality is a function of incubation temperature. In other words, the value of incubation temperature reported in the study from which the data were collected best explains mortality. For clarity we refer to this variable as “observed incubation temperature”, or *T*_*OBS*_. Second, we reasoned that TPiv and embryonic thermal tolerance exhibit a correlated evolution, in which case embryonic mortality would be best explained by the difference between TPiv and *T*_*OBS*_. To create this new variable, we subtracted *T*_*OBS*_ from species-specific TPivs, and we refer to this new variable as “deviation from TPiv”, or Δ*T*_*TPIV*_. Finally, we reasoned that if the mean incubation temperature chosen by researchers is generally similar to the mean temperature of the natural incubation environment, then the mean temperature of a given study provides a good reference point for the average incubation conditions. Mortality may then be related to the extremity of the incubation treatment and would be best explained by the difference between the mean incubation temperature of the study and *T*_*OBS*_. To create this new variable, we subtracted each value of *T*_*OBS*_ from the mean temperature of its respective study, and we refer to this new variable as “deviation from mean temperature”, or Δ*T*_*Tμ*_. We fit the three mixed linear models using maximum likelihood, where the sole fixed-effect predictor of embryonic mortality was *T*_*OBS*_ (Model A), Δ*T*_*TPIV*_ (Model B), or Δ*T*_*Tμ*_ (Model C). Models were compared using AICc. We ran these analyses twice, once with all available data, which includes all FMF and MF species (n = 199), and again after excluding FMF species, thereby including only MF species (n = 159).

We also investigated whether there is an association between sex ratio and embryonic mortality, and if so, whether this association is realized over-and-above any temperature effect. This analysis leverages the subset of data for which both temperature and sex ratio are measured at a given incubation temperature. We fit the model that best-explained variation in embryonic mortality (which was Model B, in which temperature is expressed as Δ*T*_*TPIV*_, see Results), and compared this model to two new models: Model D, in which mortality was expressed as a linear effect of sex ratio (proportion of clutch male, or *PropMale*), and Model E, in which mortality was expressed as a linear effect of *PropMale* and Δ*T*_*TPIV*_ (as in Model B). We ran separate analyses for MF species alone, then for both FMF and MF species together (FM species: 19 studies, 13 species, 130 observations; FMF & MF species: 21 studies, 14 species, 158 observations). Finally, we also fit *PropMale* as a function of Δ*T*_*TPIV*_, with StudyID as a random effect, simply to visualize how *PropMale* changes with temperature across broad species.

Theory suggests mortality should increase more rapidly with temperature under female-producing temperatures, compared to male producing temperatures (Figure 1). We therefore tested whether the relationship between mortality and temperature is different above the TPiv compared to below the TPiv. We took the absolute value of Δ*T*_*TPIV*_, thereby creating a new variable which we refer to as “absolute deviation from TPiv”, or | Δ*T*_*TPIV*_ |. We also created a categorical variable “direction of deviation”, which identifies the direction of the temperature difference with respect to TPiv (i.e., above TPiv or below TPiv). We used model selection to compare three models, and continuous fixed effects were fit as basis splines. First, we fit Model F, in which mortality was expressed simply as | Δ*T*_*TPIV*_ |. Next, we fit Model G, in which we fit |Δ*T*_*TPIV*_| and the additional term “direction of deviation”, which explores whether average mortality is different above vs below the TPiv. Finally, we fit Model H, which included an interaction between “direction of deviation” and | Δ*T*_*TPIV*_ |, which explores whether shape of the relationship between embryonic mortality and temperature differs above vs below TPiv. We ran these analyses twice, once with all available data, which includes all FMF and MF species (n = 199), and again after excluding FMF species, thereby including only MF species (n = 159).

## Results

We tested whether embryonic mortality is best explained by incubation temperature per se (*T*_*OBS*_, Model A), the difference between incubation temperature and TPiv (Δ*T*_*TPIV*_, Model B), or the difference between incubation temperature and the mean incubation temperature of the study (Δ*T*_*Tμ*_, Model C). We found no support for Model A (k = 6, LogLik = −356.4, ΔAIC = 43.3, *w*_*i*_ = 0, n = 199) or Model C (k = 6, LogLik = −353.3, ΔAIC = 37.1, *w*_*i*_ = 0, n = 199). Instead, Model B was strongly supported (k = 6, LogLik = −334.7, *w*_*i*_ = 1, n = 199). Visualization of Model B indicates that the relationship between Δ*T*_*TPIV*_ and embryonic mortality is concave-upward, centered below TPiv (Table S1, Figure 2a). We reran this analysis excluding all FMF species, and Model B remained highly supported (k = 6, LogLik = −258.7, *w*_*i*_ = 1, n = 159, Figure 2b), whereas Model A (k = 6, LogLik = −273.0, ΔAIC = 28.5, *w*_*i*_ = 0, n = 159) and Model C (k = 6, LogLik = −278.6, ΔAIC = 39.6, *w*_*i*_ = 0, n = 159) remained unsupported.

**Figure 2:**
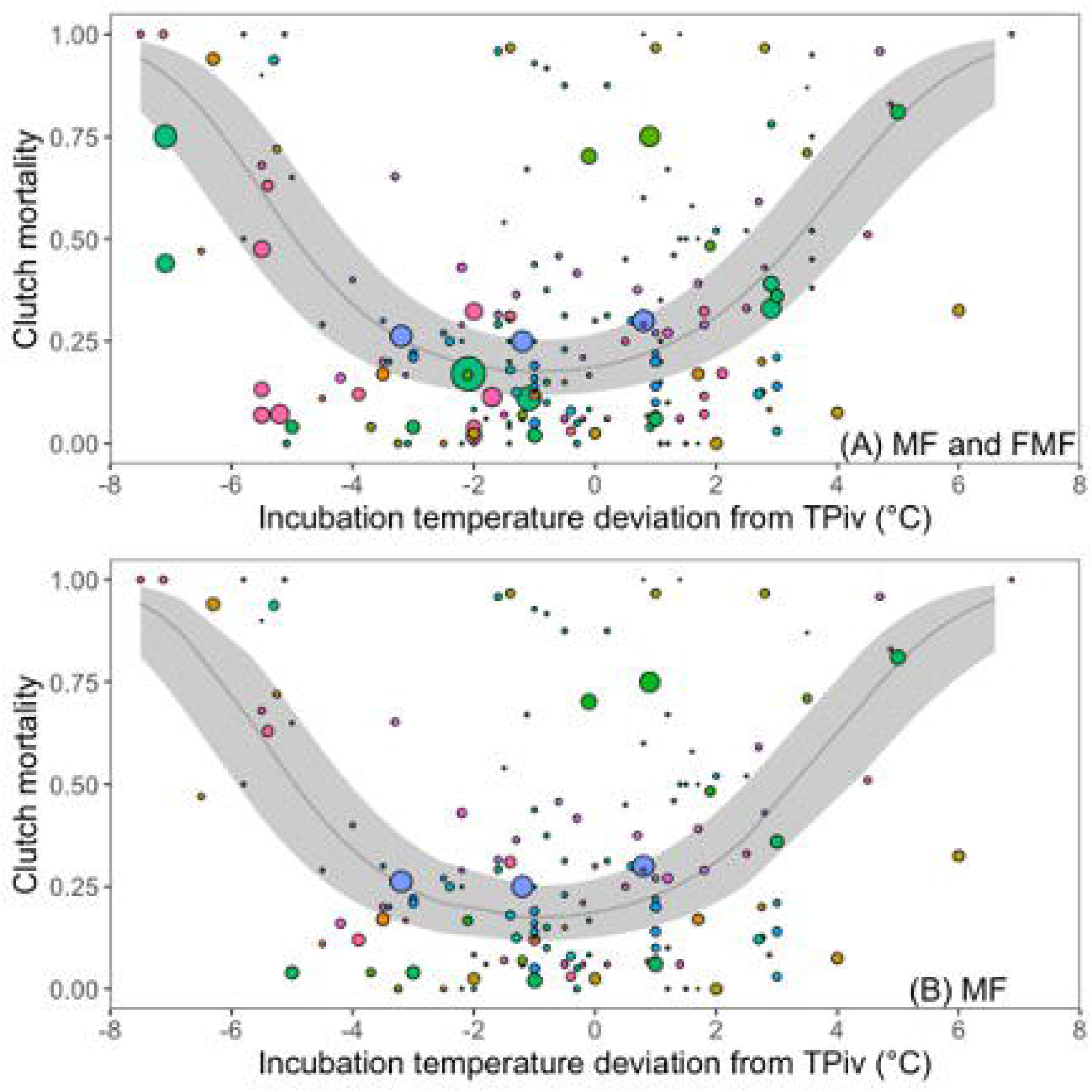
The relationship between proportion clutch mortality and incubation temperature deviation from TPiv (Δ*T*_*TPIV*_), which is the difference between incubation temperature and species-specific TPivs. In panel (a) all MF & FMF species are included, and in panel (b) FMF species are excluded. Points represent raw mortality data collected from published literature, the size of the point reflects relative sample size of each data point, and color reflects study ID. The 95% confidence intervals are in shaded grey.

Visualization of sex ratios (*PropMale*) as a function of Δ*T*_*TPIV*_ revealed the expected patterns of variation: when data from FMF and MF species are combined there is a well-defined peak of male producing temperatures below species-specific (upper) TPivs (Figure 3A), and when MF species are considered alone there was relatively little variation in sex ratios below species-specific TPiv (Figure 3B). Next, we investigated whether embryonic mortality is best predicted by temperature or sex, and we did so by modelling whether mortality was associated with *PropMale*, or Δ*T*_*TPIV*_, or both *PropMale* and Δ*T*_*TPIV*_. Of the three models compared, we found that the model featuring *PropMale* as the sole predictor of embryonic mortality, Model D, was poorly supported (k = 4, LogLik = 270.3, ΔAIC = 44.3, *w*_*i*_ = 0, n = 158). In fact, Model D was poorly supported even though there was a linear association between *PropMale* and embryonic mortality in this model (*b*_*SexRatio*_ ± *SE* = −1.02 *±* 0.210, Figure 4a). The most strongly supported model, Model B, featured Δ*T*_*TPIV*_ as the sole predictor of embryonic mortality (k = 6, LogLik = −246.0, *w*_*i*_ = 0.75, n = 158). Finally, the model featuring both *PropMale* and Δ*T*_*TPIV*_, Model E, was not supported: it differed from the best model (Model B) by one parameter and likelihoods were nearly identical (see Burnham and Anderson, 2010, pg 131) (k = 7, LogLik = −246.0, ΔAIC = 2.19, *w*_*i*_ = 0.25, n = 158). Importantly, the marginal association between *PropMale* and embryonic mortality was weak in Model E, i.e., when controlling for Δ*T*_*TPIV*_ (*b*_*SexRatio*_ = 0.0187 *±* 0.322), indicating that temperature, not sex, influences mortality. Next, we excluded all FMF data and reran the above analyses, revealing qualitatively identical results. We found that Model D was unsupported (k = 4, LogLik = −216.9, ΔAIC = 36.2, *w*_*i*_ = 0, n = 130) even though there was a non-trivial association between *PropMale* and embryonic mortality (*b*_*SexRatio*_ = −0.638 *±* 0.237, Figure 4B). Model B was again supported (k = 6, LogLik = −196.6, *w*_*i*_ = 0.72, n = 130), and the marginal association between *PropMale* and embryonic mortality was weak when controlling for Δ*T*_*TPIV*_ (*b*_*SexRatio*_ = −0.200 *±* 0.330).

**Figure 3:**
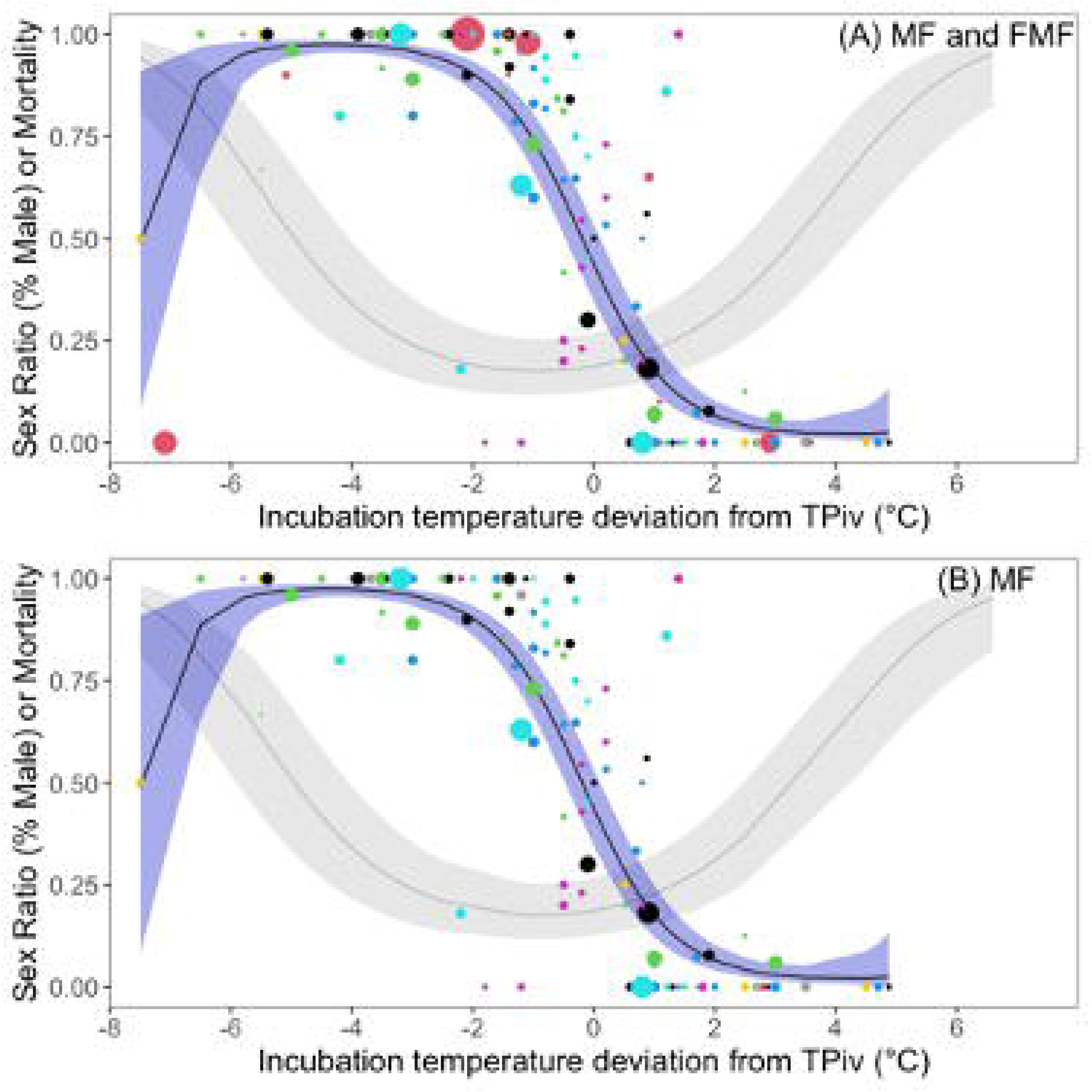
Visualization of sex ratio (proportion of clutch male) as a function of the difference between observed incubation temperature and species-specific TPivs (Δ*T*_*TPIV*_). Fitted line for sex is in blue. In grey, we overlaid the relationship between clutch mortality and Δ*T*_*TPIV*_, estimated from Model B (see Figure 2). (A) Both MF and FMF species are included in the model, and in (B) FMF species are excluded. Color of points indicate different studies and size of points indicates relative sample size. The single MF species in panel B with a prevalence of males below TPiv is the painted turtle (Schwarzkopf and Brooks, 1985).

**Figure 4:**
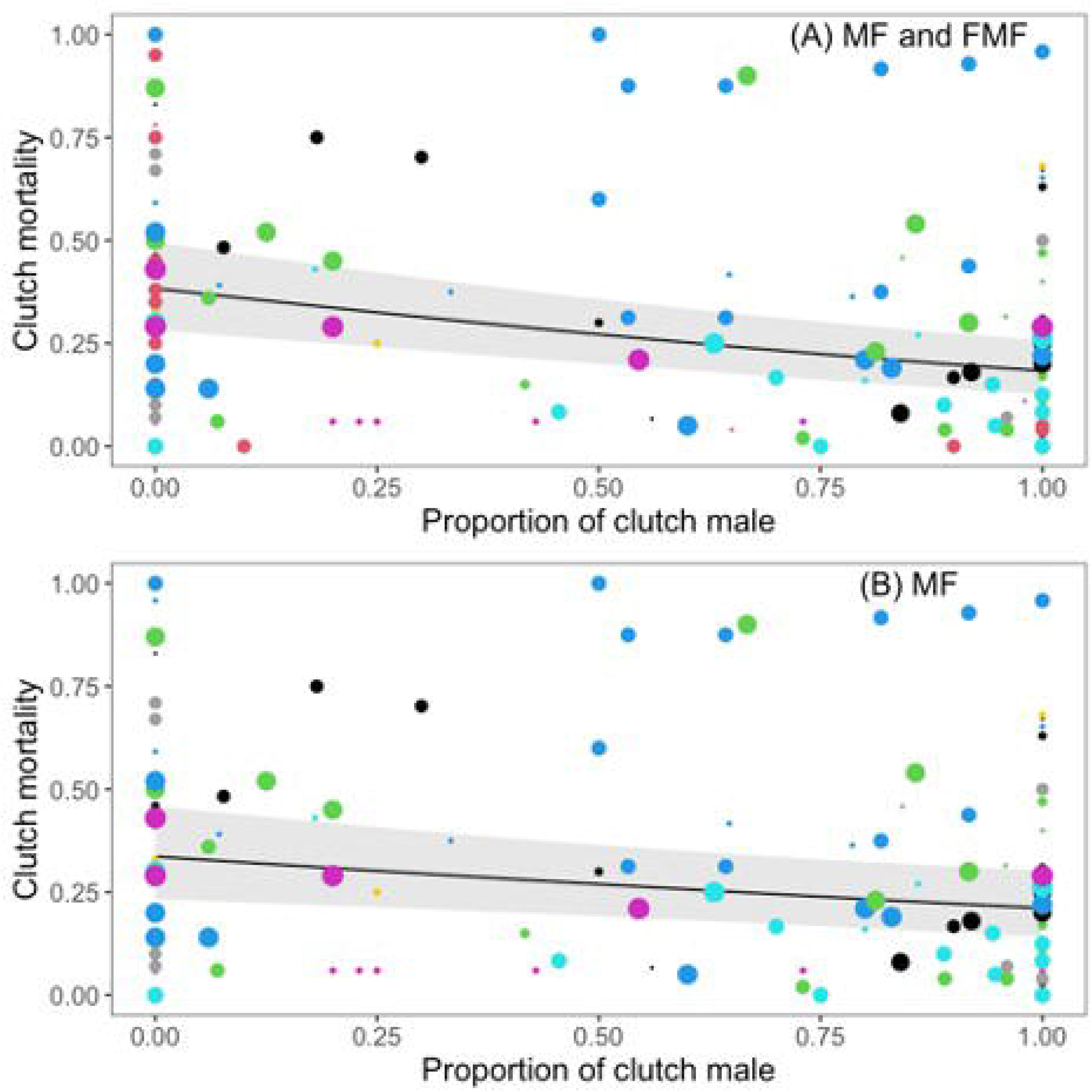
Patterns of clutch mortality as a function of proportion of clutch male for (A) FMF and MF species, and (B) MF species only. Color of points indicate different studies. Size of points reflects relative sample size of raw data points.

We tested whether patterns of temperature-dependent mortality differed above vs below species-specific TPivs. We compared three models, and we found that the model featuring an interaction between | Δ*T*_*TPIV*_ | and ‘Direction of Deviation’, Model H, was strongly supported (k = 10, LogLik = −330.7, *w*_*i*_ = 1, n = 199). The nature of the interaction in Model H supported the prediction that mortality increases relatively rapidly above the TPiv, compared to below the TPiv. Other models were poorly supported, namely Model F (k = 6, Loglik = −346.3, ΔAIC = 22.5, *w*_*i*_ = 0, n = 199) and Model G (k = 7, LogLik = −340.0, ΔAIC = 12.1, *w*_*i*_ = 0, n = 199). Next, we excluded FMF species and included only MF species in our analyses. Results were similar, as the only model without the term ‘Direction of Deviation’ was poorly supported, namely Model F (k = 6, Loglik = −261.3, ΔAIC = 6.92, *w*_*i*_ = 0.02, n = 159). Support was otherwise similar between Model G (k = 7, LogLik = −257.2, ΔAIC = 0.93, *w*_*i*_ = 0.38, n = 159) and Model H (k = 10, LogLik = −253.4, *w*_*i*_ = 0.60, n = 159, Figure 5B). Visualization again supports the interpretation that mortality increases relatively rapidly above the TPiv, compared to below the TPiv, at least for the first 5-6°C above TPiv (Figure 5B).

**Figure 5:**
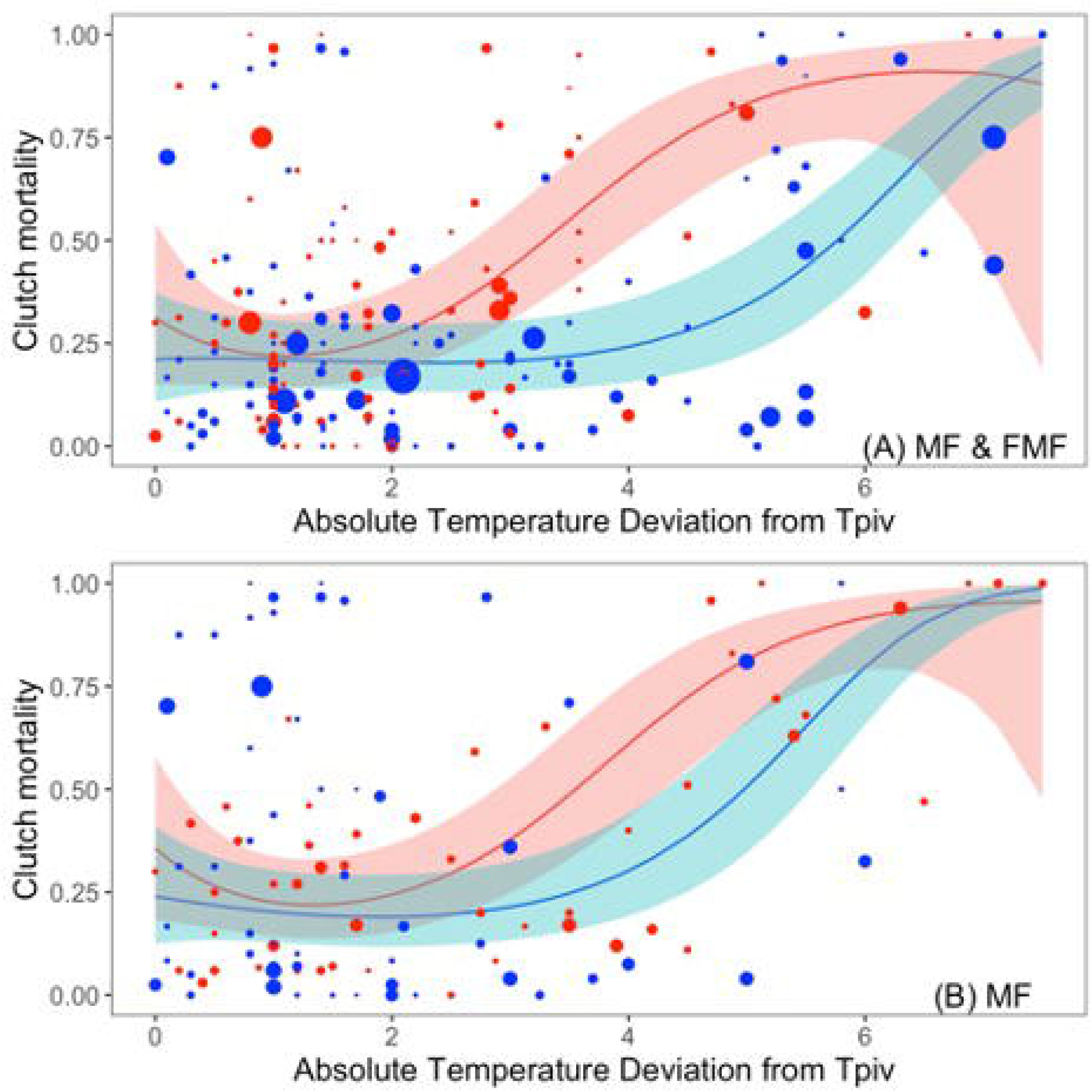
Clutch mortality as a function of the absolute difference between incubation temperature and species-specific TPivs (| Δ*T*_*TPIV*_ |) and the direction of the difference (above vs below TPiv) for (A) FMF and MF species, and (B) MF species only. Blue indicates raw data points below TPiv and red indicates raw data points above TPiv. Size of points reflects relative sample size of raw data points.

## Discussion

The present study presents a simple theory of TSD evolution (Figure 1) to explore how constant temperature incubation is associated with embryonic mortality in chelonians. We communicate four main findings. First, we show that embryonic morality is best explained by the temperature difference between species-specific TPiv and incubation temperature (Δ*T*_*TPIV*_). Second, and similarly, we find mortality is lowest near TPiv, and highest when the difference between TPiv and the incubation temperature is large. We interpret these first two patterns as consistent with the prediction that TPiv exhibits correlated evolution with embryonic thermal tolerance. Third, we found that embryonic mortality was negatively correlated with sex ratio (proportion male) for both MF species alone and for FMF and MF species combined, but that sex ratio was not associated with mortality after accounting for Δ*T*_*TPIV*_. This is consistent with the notion that temperature, not sex, exhibits an overarching effect on performance, and that female-producing thermal environments are relatively stressful for embryos. Finally, and relatedly, we demonstrate that embryonic mortality increases relatively rapidly with temperature above the TPiv, compared to below the TPiv, which is also consistent with the hypothesis that female-producing temperatures are relatively stressful for embryos. Below we explore these results and emphasize their significance in evolution and conservation.

The relatively rapid increase in mortality with temperature above the TPiv has important evolutionary implications. Mechanistically, fitness may depreciate rapidly above the TPiv because TPiv evolves to be closer to the upper end of thermal tolerance, rather than the lower end, as reaction rates for most or all processes changes more rapidly near the upper tolerance limit (Sharpe and DeMichele, 1977; Kingsolver, 2009). Indeed, the lowest level of mortality observed in the present study was just below TPiv, suggesting TPiv often falls toward the upper limit of an intermediate thermal range that is optimal for development (Stubbs and Mitchell, 2018). One interpretation of our results, therefore, is that female-producing temperatures coincide, at least roughly, with a departure from optimal incubation temperatures. Indeed, we also found that sex ratio (percent male) *per se* was negatively correlated with mortality.

Although this pattern could arise from differential mortality of male embryos when overall clutch mortality is relatively high, there is no obvious mechanism that would explain this phenomenon. We therefore suggest that the negative correlation between sex ratio (percent male) *per se* and embryonic mortality arises because female-producing temperatures are relatively stressful. Our findings are therefore important as they are consistent with the simple explanation for TSD explored herein (Figure 1). Specifically, classic theory suggests females in age-structured populations should produce more sons when sons are expected to become high-condition adults (Trivers and Willard, 1973), where condition refers to general health, energy reserves, vigor, and overall quality. Therefore, if incubation temperature has, on average, an effect on the general health and vigor of individuals at adulthood, then a sex by incubation temperature interaction for fitness (Charnov and Bull, 1977) is inexorable when males are produced under favourable incubation temperatures and females under unfavourable incubation temperatures. We suggest that this explanation for TSD should be limited to the long-lived chelonians and crocodilians, and not necessarily extended to squamates and the tuatara. This is because chelonians and crocodilians share an evolutionary history, because neither group features the FM pattern of TSD, and because the longevity of both chelonians and crocodilians results in populations that are highly age- and size-structured, where variation in male quality can be maintained.

Pivotal temperatures appear to be heritable and may be under weak selection (Beukeboom and Perrin, 2014), and there is evidence, albeit mixed, that TPivs follow broad environmental gradients in temperature (Ewert *et al.*, 2004, 2005; but see Carter *et al.*, 2019). Evolution of TPivs is therefore possible, and another prediction from the simple theory of TSD (Figure 1) is that TPiv and embryonic thermal tolerance experience a correlated evolution, as would be necessary to ensure males are produced in relatively high-quality environments. Admittedly, we define ‘tolerance’ fairly broadly, in that tolerance encompasses the range of thermal environments that support normal embryonic development, and we assume that embryonic mortality and tolerance are strongly associated.. Nevertheless, we found mortality was best explained by Δ*T*_*TPIV*_ (Model B), rather than the difference between observed temperature and the mean temperature used in the study (Δ*T*_*Tμ*_, Model C), or the observed incubation temperature (*TOBS*, Model A). One interpretation of this pattern is that TPiv and embryonic thermal tolerance undergo a correlated evolution, although the extent to which this particular finding supports our simple theory of TSD is debatable. We suggest that good evidence of a correlated evolution of TPiv and embryonic thermal tolerance would indeed arise if Δ*T*_*TPIV*_ (Model B) better-explained mortality patterns than deviation from the long-term average environmental temperature, but our variable Δ*T*_*Tμ*_ (Model C) is the average temperature used in a particular study, which is not necessarily the long-term average temperature of the local environment. In sum, it is unclear if our study provides evidence for correlated evolution between TPiv and embryonic thermal tolerance, consistent with TSD theory (Figure 1), or whether relatively strong support for our Model B vs Model C arose because researchers choose values of Δ*T*_*Tμ*_ that do not reflect the typical incubation environments of their focal populations. In any event, it is necessary to study further whether TPivs are strongly linked with thermal tolerance and physiology of the embryo, as this may provide insight into the adaptive significance of TSD.

The overarching message communicated by the present study is that a constant temperature used to produce females is associated with thermal stress, measured here as mortality. Although we did not measure post-hatching phenotypes (but see below), we argue that negative effects of thermal stress are likely to persist beyond the embryo stage. Indeed, persistent phenotypic effects of thermal stress would have ramifications for incubation techniques used in conservation programs, and such effects would be consistent with our simple model for TSD evolution. Stress is a physiological state in which normal function is disrupted, and stress may have a negative fitness impact (Walker *et al.*, 2005; Klockmann *et al.*, 2017). Our proxy of stress (mortality) represents the most extreme fitness outcome that is possible, as mortality represents an inability to persist, let alone reproduce (Kingsolver, 2009). Thermal stress can be induced at temperatures which do not necessarily induce mortality but may induce phenotypic morbidity (Cavallo *et al.*, 2015; Refsnider *et al.*, 2015; Kingsolver and Woods, 2016). Even within the present study, for example, several studies reported abnormalities in hatchlings reared in high temperature incubation trials (Packard *et al.*, 1987; Zhu *et al.*, 2006), and previous studies suggest extreme temperatures lead to developmental instability (Noble *et al.*, 2018b). We therefore suggest that our findings are probably not restricted to the effect of temperature on embryos: the relationship between temperature and fitness observed herein (i.e., embryonic mortality is lowest just below TPiv, and increases more rapidly above TPiv than below TPiv) may persist into later life by virtue of the thermal stress survivors experienced as embryos.

The form in which thermal stress might persist in post-hatching phenotypes is unclear. However, thirteen studies included in the present analysis included a measure of post-hatching growth, performance metrics, or mortality, encompassing a total of twenty-three discrete measurements (Table 1). Fifty-two percent of the measurements indicated a significant temperature effect. Due to the heterogeneity of the papers and fitness metrics, and the multi-level comparisons within some studies, a formal meta-analysis was not possible. However, our inclusion criteria for the present study likely resulted in the exclusion of many studies examining post-hatching growth or performance, such that it may prove fruitful for future research to formally examine post-hatching growth and performance under different incubation regimes, as evidence suggests the impacts of temperature stress may be observed at a variety of life stages (Jonsson *et al.*, 2014; Noble *et al.*, 2018b).

**Table 1:**
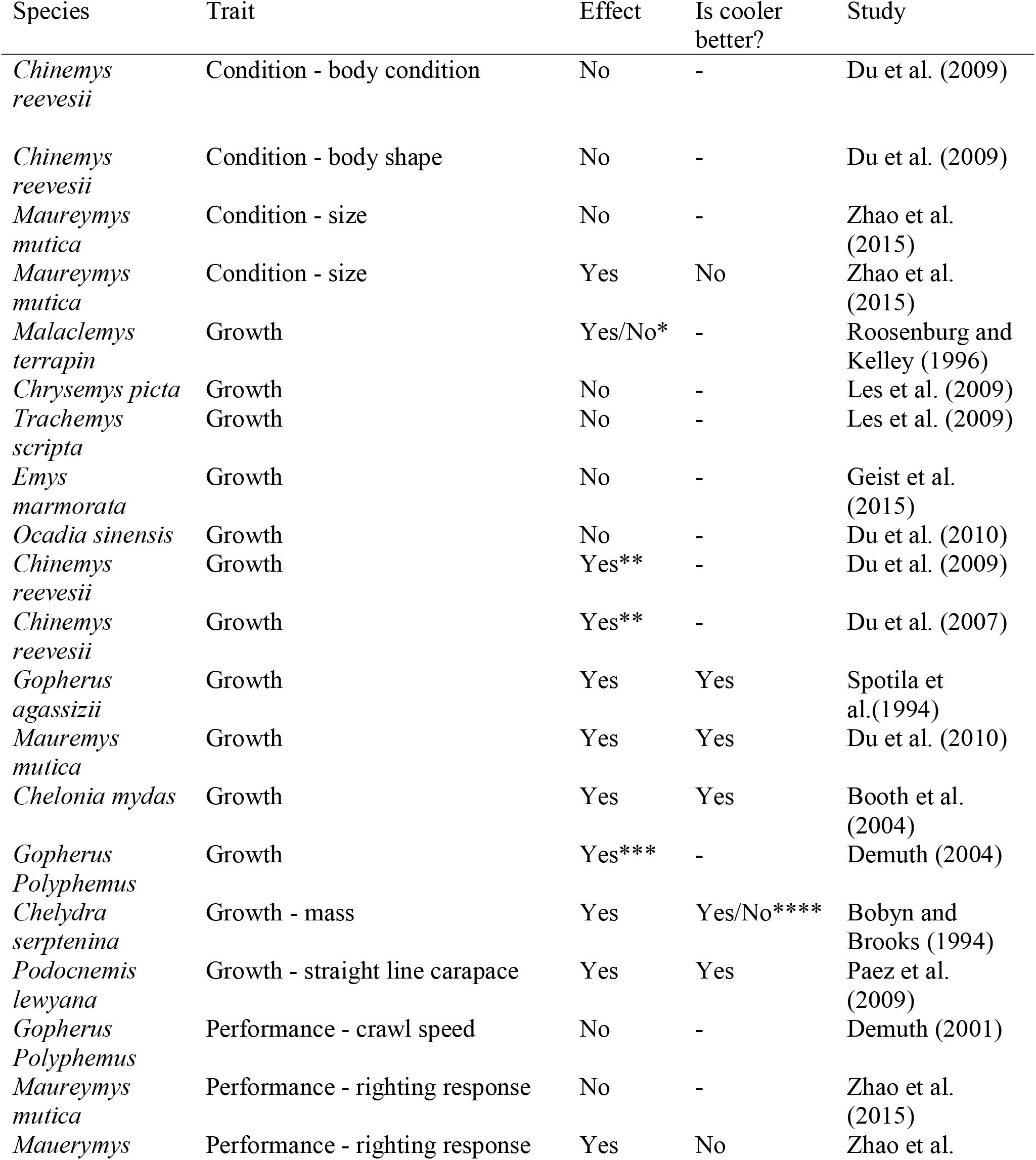
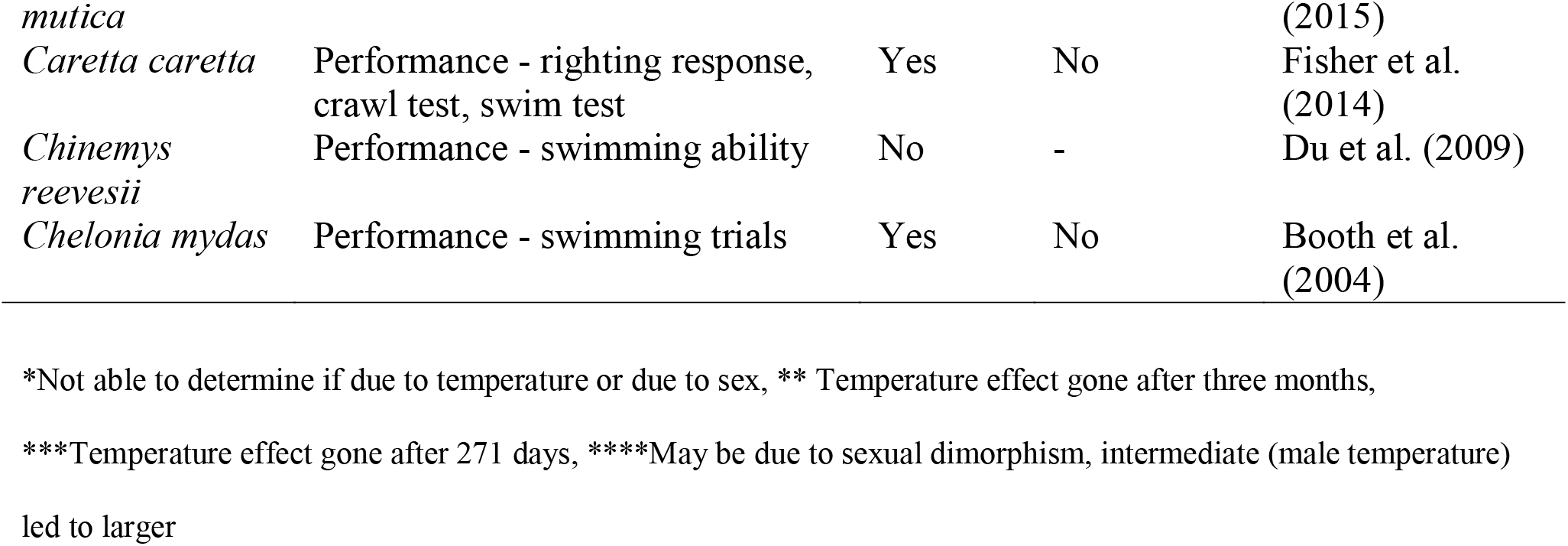
Post-hatching performance and growth temperature effect for papers included in the current study.

If artificial incubation is geared toward conservation, then presumably artificial incubation regimes fall within a normal range of incubation temperatures experienced by the focal population. However, conservation managers face at least two reasons that incentivize high-temperature incubation of turtle embryos. The first is that most turtles exhibit TSD, and for turtles, relatively warm temperatures result in an over-production of female hatchlings (Ewert *et al.*, 1994). Given that a major goal of conservation programs is ultimately to increase the number of wild births in the population, and births are more strongly related to female than male abundance (Girondot *et al.*, 2008), TSD may represent an incentive to incubate at warm temperatures. Second, high temperatures result in faster development rate and hence shorter incubation time, potentially reducing the expense associated with husbandry, further incentivizing high temperature incubation (Du *et al.*, 2006). Despite the incentives associated with high-temperature incubation, our study suggests constant temperature incubation near TPiv results in the highest hatching success and perhaps the least thermal stress. Notably, female production does not require extreme temperatures, as female-biased sex ratios can be produced at temperatures that are only slightly above the TPiv. For example, or analysis suggests that by incubating embryos only 1°C above TPiv (Figure 3), a sex ratio of over 75% females can be achieved, on average, with a trivial increase in embryonic mortality (and presumably, a trivial increase in post-hatching stress), although we note that the actual sex ratio achieved will depend on the shape of the temperature-sex reaction norm at the population level (Bentley *et al.*, 2017; Carter *et al.*, 2019). The notion of incubating embryos as near as possible to the pivotal temperature may be especially useful for head starting programs, where incubation protocols can be reviewed to prevent the release of hatchlings or juveniles that underwent significant thermal stress during development.

While some studies suggest head-starting can be beneficial (Cunnington and Brooks, 1996; Crowder and Heppell, 2011; Mullin, 2019), the long-term benefits of head starting are often unclear or negligible (Páez et al., 2015; Spencer, Van Dyke, & Thompson, 2017), as factors such as predation and adverse environmental conditions result in low natural or even negligible survival of hatchlings and juveniles in wild environments (Heppell, 1998; Enneson and Litzgus, 2008; Altobelli, 2017). Although the present study does not definitively link incubation environments with long-term survival as we do not examine post-hatching fates, we suggest that incubation of embryos near TPiv may ultimately improve survival prospects of individuals, especially females, released into the wild. As we have pointed out, incubation near TPiv is compatible with sex ratio manipulation for many species and populations with TSD. Indeed, it is desirable when increases in adult recruitment translate into increases in juvenile recruitment, and this requires that careful attention is paid to the sex of released individuals, as population size is not indicative of extinction risk when sex ratios are skewed (Dale, 2001; Clout *et al.*, 2002; Wedekind, 2002; López-Sepulcre *et al.*, 2009).

Data syntheses and analyses are able to identify large-scale patterns, and ultimately help strengthen evidence-based decision-making (Stewart, 2010). Indeed, it is useful for conservation managers to periodically review methodologies used for species recovery, and with the substantial increase in reptile conservation organizations since the 1990s, there are likely many different initiatives that leverage artificial incubation (Gibbons and Lovich, 2019). The present study underlines how constant temperature incubation in artificial egg incubation regimes affects embryonic mortality, and possibly post-hatching thermal stress. We emphasize that we studied chelonians in particular, but similar studies of crocodilians, and especially squamates and the tuatara where different patterns of TSD can be found (Valenzuela and Lance 2004), would be useful to uncover how TPiv relates to mortality in those groups. More broadly, we suggest chelonian conservation programs leveraging artificial incubation may benefit from avoiding high temperature incubation environments, and incubating embryos near species-specific TPivs.

## Acknowledgements

We thank Anthony Rajkumar, Sami Troendle for constructive comments on the original manuscript, and Melanie Massey for help compiling data. We also thank Daniel Noble and two anonymous reviewers for their generous, constructive, and extensive feedback that greatly improved the quality of this work. Funding was provided by an NSERC Discovery Grant Number 2016-06469 to NR.

